# Targeting Wnt Signaling To Overcome PARP Inhibitor Resistance

**DOI:** 10.1101/378463

**Authors:** Tomomi M. Yamamoto, Alexandra McMellen, Zachary L. Watson, Jennifer Aguilera, Matthew J. Sikora, Rebecca Ferguson, Elmar Nurmemmedov, Tanay Thakar, George-Lucian Moldovan, Hyunmin Kim, Diana M. Cittelly, Heidi Wilson, Kian Behbakht, Benjamin G. Bitler

## Abstract

Epithelial ovarian cancer (EOC) has one of the highest deaths to incidence ratios. High grade serous ovarian carcinoma (HGSOC) is the most common and deadliest EOC histotype because of the lack of secondary therapeutic options following debulking surgery and platinum/taxane-based chemotherapies. For recurrent chemosensitive HGSOC, poly(ADP)-ribose polymerase inhibitors (PARPi; olaparib, rucaparib, or niraparib) represent an emerging treatment strategy. While PARPi are most effective in homologous recombination DNA repair-deficient (HRD) HGSOCs, more recent studies have observed a significant clinical benefit in non-HRD HGSOCs. However, all HGSOC patients are likely to acquire resistance to PARPi. Therefore, there is an urgent clinical need to better understand PARPi resistance, and to introduce novel combinatorial therapies to overcome PARPi resistance and extend HGSOC disease-free intervals. Utilizing a two *BRCA2-mutated* and one BRCA-wildtype HGSOC cell lines that are olaparib sensitive, we established resistant cells. Transcriptome analysis of the matched olaparib-sensitive versus resistant cells did not detect *BRCA2* reversion mutations, but revealed activation of Wnt/TCF signaling pathway, as TCF transcriptional activity was significantly increased in PARPi-resistant cells. In parallel, forced activation of Wnt signaling in PARPi-sensitive cells via WNT3A stimulation reduced response to PARPi. In a recurrent-HGSOC PARPi insensitive patient-derived xenograft model there was an increase in a Wnt/TCF transcriptional target. PARPi resistant cells were sensitive to inhibition of Wnt signaling using the FDA-approved compound, pyrvinium pamoate, which has been shown to inhibit Wnt signaling. We observed that combining pyrvinium pamoate with olaparib resulted in a significant decrease in tumor burden and number of tumor nodules. This study demonstrates that Wnt signaling can mediate PARPi resistance in HGSOC and provides a clinical rationale for combining PARPi and Wnt inhibitors.

## INTRODUCTION

Epithelial ovarian cancer (EOC) is the deadliest gynecological malignancy and has one of the highest deaths to incidence ratios (63 deaths:100 cases) (1). The most common EOC histotype is high grade serous ovarian carcinoma (HGSOC) and over 85% of HGSOC are diagnosed at late stages (III/IV) (1). About 80% of all HGSOC patients respond to first-line therapy, which includes surgical debulking and platinum/taxane-based chemotherapies. However, HGSOC recurs in a majority of patients, and patients are subsequently treated with additional chemotherapeutic regimens (2). As tumors acquire chemoresistance, disease-free intervals are shortened with each subsequent recurrence, which makes identifying effective and novel therapeutic strategies for recurrent HGSOC an urgent clinical need.

The Cancer Genome Atlas revealed approximately 50% of all HGSOC tumors have mutations or deficiencies in the homologous recombination (HR) DNA repair pathway (3). For example, mutations and epigenetic silencing of *BRCA1/2* are detected in 30% of HGSOC cases. HR-deficient cancers can be targeted in a synthetic lethal fashion using poly(ADP)-ribose polymerase (PARP) inhibitors (2,4–6). There are currently three PARP inhibitors (PARPi; olaparib, rucaparib, and niraparib) that are FDA-approved for the treatment of recurrent HGSOC. Patients with HR-deficient HGSOC tumors treated with PARPi have a significant clinical benefit but recent clinical trials have observed a clinical benefit in patients without measurable HR-deficiency (7,8). Nevertheless, in a similar fashion as first-line chemotherapies, PARPi treated patients will likely recur with resistant disease.

Tumorigenesis and chemoresistance in numerous cancer types, including ovarian cancer can be attributed to Wnt signaling. Canonical Wnt signaling is mediated through ligand (i.e. WNT3A) stimulation of frizzled (FZD) and lipoprotein receptor-related protein (LRP) receptors. WNT-receptor interactions promote sequestration of the β-catenin degradation complex and accumulation and nuclear localization of β-catenin. Increased β-catenin leads to its interaction with T-cell factor (TCF) and lymphoid enhancer factor (LEF) transcriptional activators and ultimately upregulation of TCF/LEF target genes (e.g. *FOSL1*). Previously, we established in HGSOC cell lines β-catenin-dependent TCF transcriptional activation is antagonized by non-canonical Wnt (β-catenin-independent) signaling, and in HGSOC primary tumor the attenuation of non-canonical Wnt signaling was associated with worse overall survival (9). The observed overall survival was potentially due to the decreased time to chemoresistance. Hyperactivation of Wnt signaling has been attributed to chemotherapeutic resistance in a variety of epithelial-derived cancers including, ovarian, breast, and bladder (10–12). Based on these observations, we investigated how aberrant activation of canonical β-catenin-dependent Wnt signaling regulates PARPi response and/or resistance.

In this study, we examined HGSOC models of acquired PARPi resistance. We observed in these models that hyperactivation of Wnt signaling pathway was necessary and sufficient for PARPi resistance. In two PARPi-resistant HGSOC cell lines we observed an increase in TCF transcriptional activity compared to parental cells. We screened a panel of Wnt signaling inhibitors and observed an FDA-approved compound (pyrvinium pamoate) induced cell death in PARPi-resistant cells at nanomolar concentrations. PARPi resistant cells had increased DNA repair capacity by both non-homologous end-joining (NHEJ) and HR, independent of BRCA2 reversion mutations. Pyrvinium pamoate effectively inhibited both NHEJ and HR-mediated repair in PARPi-resistant cells, and synergized with olaparib to induce apoptosis. In an *ex vivo* culture of a primary HGSOC tumor PARPi and pyrvinium pamoate reduced proliferation. Utilizing a HGSOC cell line xenograft mouse model we demonstrated that combining olaparib with pyrvinium pamoate significantly inhibited the rate of tumor growth, disseminations and overall tumor burden *in vivo*.

## RESULTS

PARPi are most effective in homologous recombination deficient (HRD) tumors, however PARPi do convey a significant clinical benefit in most HGSOC patients (7,8). Acquired PARPi resistance needs to be examined to better understand mechanisms of action. Therefore, we established a panel of olaparib resistant HGSOC cell lines (PEO1-olaparib resistant, PEO1-OR; OVCAR10-olaparib resistant, OVCAR10-OR, and OVSAHO-olaparib resistant, OVSAHO-OR) through step-wise dose escalation of olaparib. Olaparib resistance was confirmed with a dose response colony formation assay (Fig. 1A and Sup. Fig. 1A). In PEO1-OR cells we isolated four clonal populations and confirmed >25-fold increase in half maximal inhibitory concentrations in all four PEO1-OR clones (Fig. 1B). BRCA reversion mutations are a published mechanism of PARPi resistance (13), using a N-terminal BRCA2 antibody we confirmed that BRCA2 protein expression was not restored in PEO1-OR clonal populations (Fig. 1C), suggesting an alternative mechanism of acquired olaparib resistance. Transcriptome analysis (RNA-sequencing; RNA-seq) of the PEO1-OR clones was performed to identify differentially regulated gene expression and pathways compared to olaparib-sensitive PEO1. Principal component analysis and hierarchical clustering of sequencing showed that the sensitive and resistant cells clustered together (Sup. Fig 1B-C). Although the PEO1-OR clones were independently isolated their transcriptome profiles were similar. Comparing sensitive cells to the four PEO1-OR clones there were 1,819 genes that were differentially regulated (FDR<15%, p<0.00001; Sup. Table 1). Gene Set Enrichment Analysis, pathway enrichment (KEGG) and transcription factor analysis (14,15) of differentially expressed genes was performed to identify putative pathways mediating olaparib resistance. We identified that a majority of differentially expressed genes are regulated by TCF3 and LEF1 transcription factor (Table 1). KEGG pathway analysis revealed several highly enriched pathways including, MAPK signaling, focal adhesion, and Wnt signaling (Table 2). Enrichment of TCF3/LEF1 and Wnt signaling is consistent with activation of canonical, β-catenin-dependent Wnt signaling in PEO1-OR clones. To confirm this, we examined TCF mediated transcriptional activation through a classic TCF reporter assay (TOP/FOP-FLASH), and observed a significant increase in TOP-FLASH activity in PEO1-OR and OVCAR10-OR cells versus the parental cells (Fig. 1D and Sup. Fig. 1D). OVSAHO-OR did not demonstrate a significant increase in TCF transcriptional activity (Sup. Fig. 1D). In PEO1-OR we validated several genes identified by RNA-Seq that are associated with either activation (*FOSL1, CCND1, WNT3A*) or inhibition (*WNT5A, WNT7B, SFRP1*) of canonical Wnt signaling. We confirmed that Wnt signaling activators are upregulated and Wnt signaling inhibitors are repressed in PEO1-OR cell (Fig. 1E). Notably, we observed a significant increase in *WNT3A*, a potent driver of canonical Wnt signaling. These data confirm the Wnt pathway is activated in olaparib resistant cells therefore we more closely examined the role of Wnt signaling in olaparib resistance.

**Figure 1.**
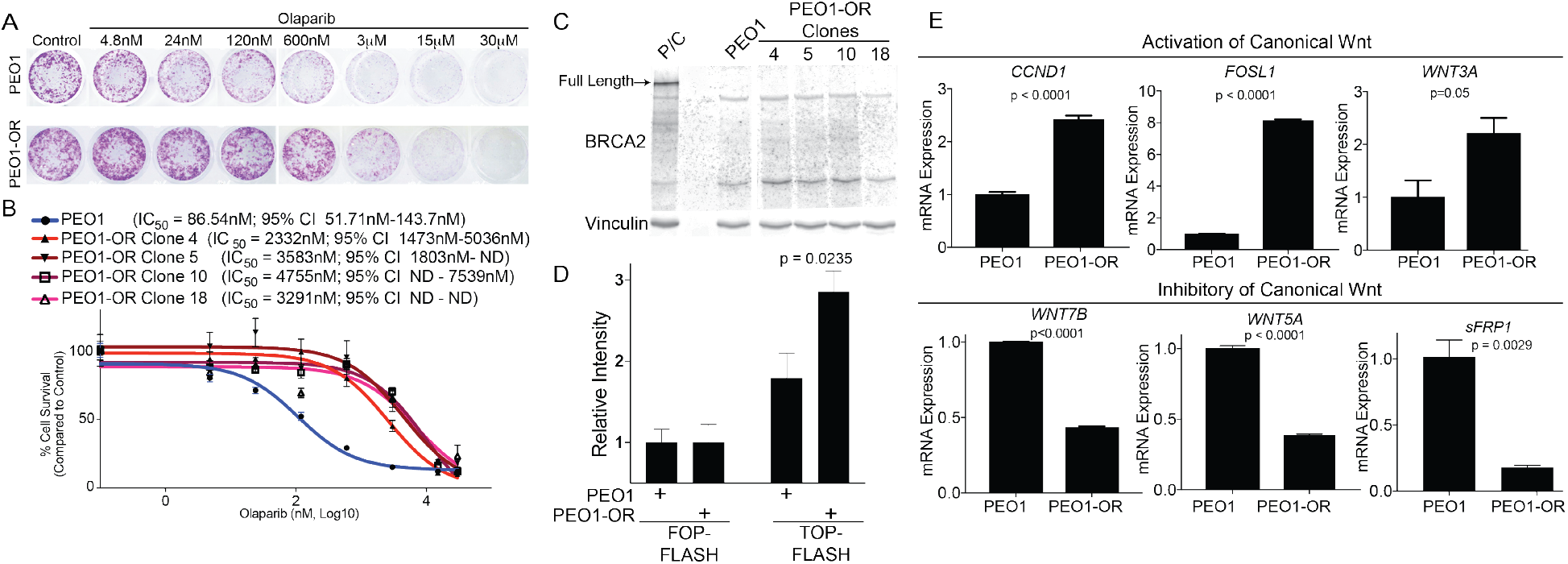
Olaparib resistant high grade serous ovarian cancer cells have increased Wnt activation. **A**) PEO1 (*TP53* and BRCA2-mutated) were treated in a step-wise fashion with increasing doses of olaparib. PEO1 sensitive and resistant (PEO1-OR) cells were plated in a 24-well plate and treated with increasing doses of olaparib for 12 days. Cells were stained with crystal violet. **B**) Four olaparib resistant populations were established and olaparib resistance was confirmed with a dose response colony formation assay. Dose response curves are graphed and IC50 with 95% confidence interval indicated. **C**) Protein was collected from PEO1 sensitive and four clonal resistant populations. Protein was separated on an SDS-PAGE and blotted against the N-terminus of BRCA2. **D**) PEO1 and PEO1-OR were transfected with TCF transcriptional reporter (TOP-FLASH) or a control reporter (FOP-FLASH). Luciferase activity was measured. **E**) RNA from PEO1 and PEO1-OR was extracted and utilized for a quantitative reverse-transcriptase PCR (qRT-PCR) against *CCND1, FOSL1, WNT3A, WNT7B, WNT5A*, and *SFRP1*. Beta-2-microglobulin (*B2M*) was used as an internal control. Experiments performed in triplicate. Statistical used to calculate p-values, unpaired two-tailed t-test. Error bars = SEM.

**Table 1.** Transcription factor enrichment in 1,819 PEO1-OR associated differentially regulated genes.

| Rank | Transcription Factor | # Genes in Gene Set | # Genes in Overlap | % of Overlap | p-value | FDR q-value |
| --- | --- | --- | --- | --- | --- | --- |
| 1 | TCF3 | 2485 | 339 | 13.64% | 8.48E-100 | 5.22E-97 |
| 2 | SP1 | 2940 | 361 | 12.28% | 3.07E-93 | 9.44E-91 |
| 3 | MAZ | 2274 | 293 | 12.88% | 1.99E-79 | 4.09E-77 |
| 4 | LEF1 | 1972 | 255 | 12.93% | 6.04E-69 | 9.29E-67 |
| 5 | UNKNOWN | 1890 | 244 | 12.91% | 1.04E-65 | 1.28E-63 |
| 6 | NFAT | 1896 | 221 | 11.66% | 1.31E-51 | 1.34E-49 |
| 7 | AP4 | 1524 | 189 | 12.40% | 6.21E-48 | 5.45E-46 |
| 8 | FOXO4 | 2061 | 223 | 10.82% | 1.15E-46 | 8.84E-45 |
| 9 | TATA1 | 1296 | 158 | 12.19% | 4.07E-39 | 2.78E-37 |
| 10 | AP1 | 1121 | 142 | 12.67% | 5.00E-37 | 3.08E-35 |

**Table 2.** KEGG pathway analysis from 1,819 PEO1-OR associated differentially regulated genes.

| Rank | KEGG | # Genes in Gene Se | # Genes in Overlap | % of Overlap | p-value | FDR q-value |
| --- | --- | --- | --- | --- | --- | --- |
| 1 | Pathways In Cancer | 328 | 62 | 18.90% | 6.12E-26 | 1.14E-23 |
| 2 | Focal Adhesion | 201 | 41 | 20.40% | 6.90E-19 | 6.42E-17 |
| 3 | Axon Guidance | 129 | 29 | 22.48% | 5.97E-15 | 3.70E-13 |
| 4 | Cell Adhesion Molecules Cams | 134 | 29 | 21.64% | 1.72E-14 | 8.01E-13 |
| 5 | MAPK Signaling Pathway | 267 | 41 | 15.36% | 2.22E-14 | 8.27E-13 |
| 6 | Regulation Of Actin Cytoskeleton | 216 | 35 | 16.20% | 3.46E-13 | 1.07E-11 |
| 7 | Leukocyte Transendothelial Migration | 118 | 25 | 21.19% | 1.80E-12 | 4.78E-11 |
| 8 | WNT Signaling Pathway | 151 | 28 | 18.54% | 2.72E-12 | 6.32E-11 |
| 9 | T Cell Receptor Signaling Pathway | 108 | 23 | 21.30% | 1.23E-11 | 2.53E-10 |
| 10 | Small Cell Lung Cancer | 84 | 19 | 22.62% | 2.45E-10 | 4.55E-09 |

PARPi resistant cells have increased Wnt signaling, therefore we wanted to know if Wnt hyperactivation was sufficient to drive PARPi resistance. To examine the impact on increased Wnt/TCF activation on olaparib response, the open-reading frame of a Wnt ligand (WNT3A) was transduced into olaparib-sensitive PEO1 cells. WNT3A expression was confirmed to be significantly upregulated through quantitative PCR (qPCR) and immunoblot (Fig. 2A-B). In PEO1-WNT3A, TCF transcriptional activity was significantly increased (Fig. 2C), and canonical Wnt target genes (*CCND1* and *FOSL1*) were upregulated following WNT3A overexpression (Fig. 2D). Next, olaparib sensitivity was assessed in PEO1-WNT3A via dose response colony formation. The overexpression of WNT3A promoted olaparib insensitivity (Fig. 2E-F). The Wnt-dependent decrease in olaparib sensitivity suggests that increased Wnt signaling contributes to PARPi resistance.

**Figure 2.**
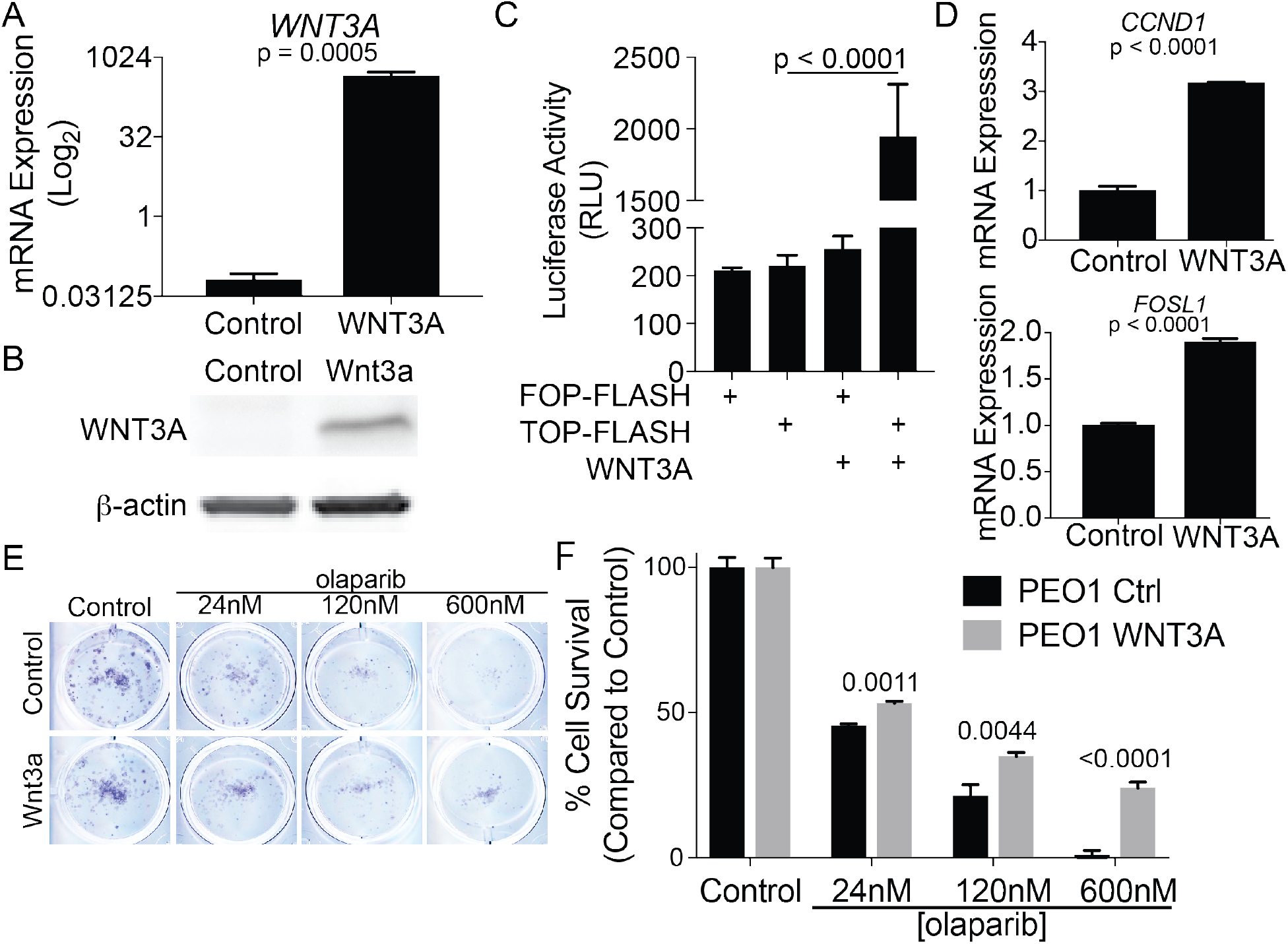
Activation of Wnt signaling contributes to olaparib resistance. **A**) PEO1 cells were transduced with either lentiviral control (Control) or a construct specific for WNT3A. mRNA expression of *WNT3A* was evaluated in PEO1-Control and PEO1-WNT3A. Beta-2-microglobulin (*B2M*) was used as an internal control. **B**) Same as A, but protein was extracted, separated on a SDS-PAGE, and immunoblotted against WNT3A. Loading control = β-actin. **C**) PEO1 control and PEO1-WNT3A cells were transfected with a TCF reporter (TOP-FLASH) and control (FOP-FLASH). **D**) RNA was extracted from PEO1 control and PEO1-WNT3A cells and used for qRT-PCR against *FOSL1* and *CCND1. B2M* was used as an internal control. **E**) PEO1 control and PEO1-WNT3A cells were plated in a 24-well plate, treated with increasing dose of olaparib for 12 days, and remaining cells were stained with crystal violet. **F**) Same as E, quantification of crystal violet staining. Experiments performed in triplicate. Statistical used to calculate p-values, unpaired two-tailed t-test. Error bars = SEM.

Since Wnt signaling was activated in olaparib-resistant HGSOC cells, and Wnt ligand over-expression was sufficient to reduce HGSOC cell sensitivity to olaparib, we examined the impact of pharmacologic inhibition of Wnt signaling when combined with olaparib. We blocked Wnt signaling by targeting distinct components of Wnt signaling, using a porcupine inhibitor (WNT-C59) to block Wnt ligand secretion, and two β-catenin inhibitors (PRI724 and Pyrvinium pamoate). PRI724 blocks β-catenin transcriptional activation by steric inhibiton of β-catenin and one if its co-activator’s CBP (CREB binding protein). Pyrvinium pamoate (Pyr. Pam.) promotes β-catenin downregulation by activating the β-catenin degradation complex in a casein kinase 1-dependent fashion (16). WNT-C59 did not reduce PEO1 cell viability at the highest concentration tested (50μM) (Fig. 3A). A secondary cell line, OVSAHO, was also utilized to determine the impact of Wnt inhibitor on HGSOC viability (Sup. Fig. 2A). Pyrvinium pamoate (Pyr. Pam.) significantly inhibited cellular viability in both PEO1 (IC50=370.2nM) and OVSAHO (IC50=733.2nM). Notably, the PEO1-OR cells were significantly more sensitive to Pyr. Pam. compared to PEO1 (IC50=169.5nM versus IC50= 315nM, p<0.0001) (Fig. 3B). OVSAHO-OR cells also were more sensitive to Pyr. Pam. (Sup. Fig. 2B) even in the absence of a significant increase in TCF activity (Sup. Fig. 1D). Taken together these data suggests an increased dependence on Wnt/β-catenin signaling in olaparib resistant cells.

**Figure 3.**
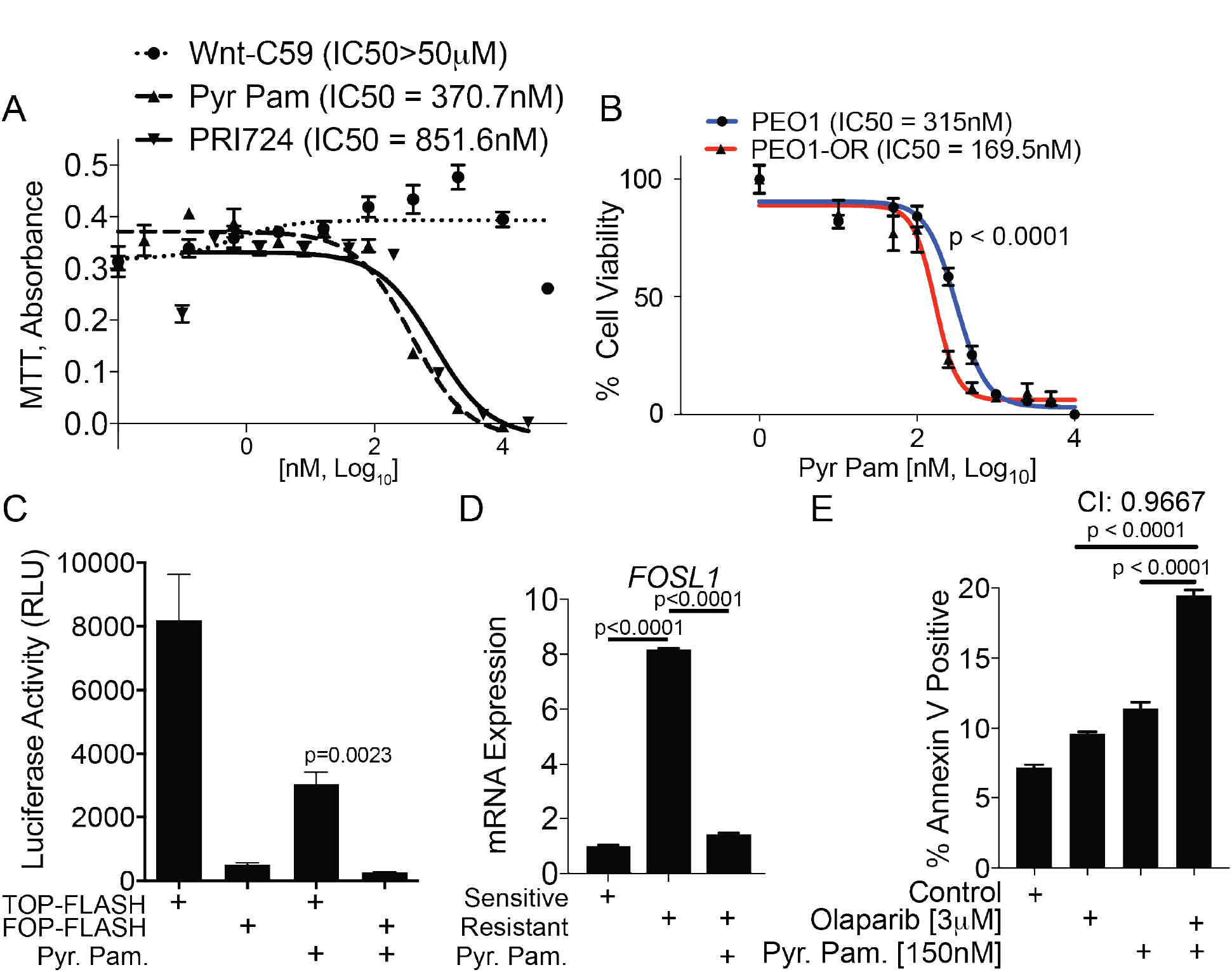
Pyrvinium pamoate significantly inhibits cell viability and synergizes with olaparib to induce apoptosis in PEO1-OR cells. **A**) PEO1 cells were treated with increasing doses of three different Wnt inhibitors (Wnt-C59, Pyr. Pam., and PRI724). Cells were treated for 48 hrs and a MTT assay was used to examine changes in cell viability. IC50 values indicated for each inhibitor. **B**) PEO1 and PEO1-OR were incubated for 48 hrs with increasing doses of Pyr. Pam. and a MTT assay was used to examine changes in cell viability. IC50 values indicated. **C**) PEO1-WNT3A cells were transfected with a TCF reporter (TOP-FLASH) and control (FOP-FLASH). Cells were treated with Pyr. Pam. (370 nM) and luciferase activity was measured. **D**) PEO1-OR cells were treated with vehicle control or Pyr. Pam. (370 nM) and RNA was extracted. RT-qPCR was performed against *FOSL1. GAPDH* was used as an internal control. P-values calculated with ANOVA. **E**) PEO1-OR cells were treated with vehicle control (DMSO), olaparib (3 μM), Pyr. Pam. (150 nM), or in combination. Cells were treated for 72 hours and subjected to an apoptosis assay (AnnexinV/PI). A combination index was calculated using CompuSyn. Experiments performed in triplicate. Unless noted, statistical used to calculate p-values, unpaired two tailed t-test. Error bars = SEM.

Pyr. Pam. is an FDA approved anthelminthic and inhibits Wnt signaling through activation of casein kinase 1 and stabilization of the β-catenin degradation complex. *In vitro* Pyr. Pam. has been reported to inhibit colorectal, ovarian and aggressive breast cancers (17–19). We confirmed that Pyr. Pam. significantly reduced TCF transcriptional activity (Fig. 3C) and repressed *FOSL1* expression in PEO1-OR cells (Fig. 3D). In addition, apoptosis was measured via Annexin/PI in PEO1-OR cells treated with a combination of olaparib and Pyr. Pam.. Examination of the single agent treatment confirmed that the PEO1-OR cells were resistant to olaparib and sensitive to Pyr. Pam. (Fig. 3E). Combining olaparib and Pyr. Pam. synergized (combination index: 0.9667) to promote apoptosis. This data suggest that treating recurrent HGSOC either with a single agent (Pyr. Pam.) or in combination (olaparib and Pyr. Pam.) could potentially promote increased tumor cell apoptosis.

We next wanted to examine the mechanism of Wnt signaling mediated PARPi insensitivity. PARPi resistance independent of a BRCA-reversion mutation has been attributed to the restoration of DNA replication fork stability (20,21) and/or increased DNA repair capacity. Given ectopic expression of WNT3A resulted in decreased olaparib sensitivity and the potential of other confounding factors in PEO1-OR cells, we examined replication fork stability following WNT3A overexpression in the sensitive PEO1 cells. PEO1 and PEO1-WNT3A cells were pulsed with 2’-deoxy-5-iodouridine (IdU), treated with hydroxyurea (HU) to promote replication fork stalling, washed, and pulsed with 5-chloro-2’-deoxyuridine (CIdU). To assess replication fork degradation while excluding premature replication termination events, lengths of IdU tracks occurring adjacent to CldU tracks were measured. Although significant, WNT3A overexpression only marginally rescued HU-induced replication fork (RF) degradation (9.21 vs. 10.43 microns for PEO1 and PEO1-WNT3A, respectively, Sup. Fig 3A). We therefore wanted to examine DNA repair capacity through functional repair assays.

We next evaluated DNA damage response by examining γH2Ax resolution and by utilizing two-plasmid functional assays. In a similar fashion as previously demonstrated (22) we utilized a marker of DNA damage (Serine 139 phosphorylated histone H2x, γH2Ax) and examined the rate of γH2Ax resolution as a DNA repair read-out. PEO1 sensitive and PEO1-OR cells were irradiated (5Gy) and γH2Ax was examined. Over an 8 hour time course PEO1-OR cells resolved γH2Ax 2.5X faster than PEO1 sensitive cells (Fig. 4A-B). We subsequently employed a functional two-plasmid DNA repair system to assess specific DNA repair pathways, HR, distal NHEJ and microhomology NHEJ (mh-NHEJ). Briefly, a unique restriction enzyme, I-SceI introduces DNA double strand breaks in a unique GFP-mutated plasmid and DNA repair by the specific pathway leads to the restoration of a GFP open-reading frame (23). To limit heterogeneity, within the assay PEO1-OR clonal populations were analyzed for the three different DNA repair pathways. Cells were incubated for 72hrs after transfection with I-SceI and using flow cytometry the percentage of GFP-positive cells was measured. Only one of the four PEO1-OR clones had a significant increase in mh-NHEJ-mediated repair compared to the sensitive control (Sup. Fig. 3B). In contrast, all four PEO1-OR clones had a significant increase in distal-NHEJ (Fig. 4C-D) and three of the four PEO1-OR clones had a significant increase in HR-mediated repair compared to the sensitive cells (Fig. 4E). Both distal-NHEJ and HR-mediated DNA repair were significantly inhibited following Pyr. Pam. treatment (Fig. 4F-G). In PEO1-OR, DNA repair activity was abrogated by β-catenin inhibition suggesting a Wnt-dependent regulatory role in conveying increased DNA repair and PARPi resistance.

**Figure 4.**
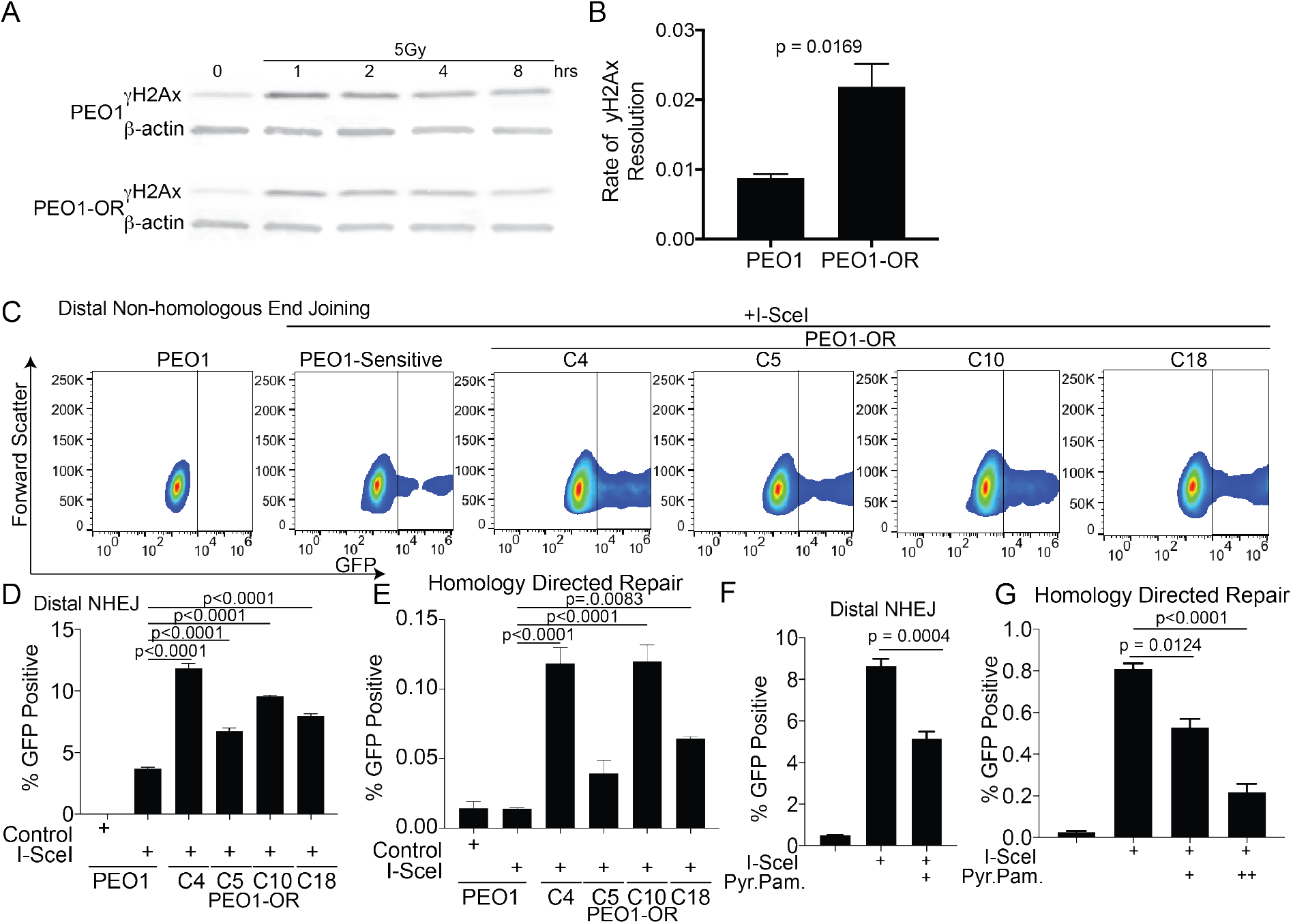
PEO1-OR have increased DNA repair capacity, which is inhibited by Pyr. Pam. **A**) PEO1 sensitive and PEO1-OR (resistant) cells were irradiated (5Gy) and incubated for the indicated time. Protein was extracted, separated with a SDS-PAGE and immunoblotted against of serine 139 phosphorylated histone H2x (γH2Ax). Loading control = β-actin. **B**) Densitometric analysis of immunoblots. Relative γH2Ax levels calculated based on β-actin levels graphed over time analyzed with a linear regression. Slope is graphed with standard error. **C**) Two-plasmid functional assay performed to assess distal non-homologous end joining. PEO1 sensitive and PEO1-OR clones were stably transfected with pimEJ5GFP and subsequently transfected with I-SceI restriction enzyme. After 72 hours transfected cells were collected and examined via a flow cytometer to quantify GFP positive cells. Representative gating strategy shown. **D**) Same as C, Distal NHEJ GFP positive cells measured for PEO1 and PEO1-OR clones. P-values calculated with ANOVA. **E**) Same as D, but utilized DRGRP to measure homology directed repair in PEO1 and PEO1-OR clones. P-values calculated with ANOVA. **F**) Same as D, but cells treated with Pyr. Pam. 24hrs after I-SceI transfection. **G**) Same as E, but treated with Pyr. Pam 24hrs after I-SceI transfection. Experiments performed in triplicate. Unless noted statistical used to calculate p-values, unpaired two tailed t-test. Error bars = SEM.

PARPi were initially developed to exploit HR-deficient tumors and restoration of HR decreases PARPi sensitivity. Therefore, we wanted to further examine HR-mediated repair through Rad51 loading onto DNA, which is a functional readout of HR (24). PEO1-OR clones were irradiated (IR, 5Gy) and incubated for 4 hours. Cells were fixed and utilized for immunofluorescence against Rad51. IR-induced Rad51-positive cells were quantified for each PEO1-OR clone and we found a significant increase in IR-induced Rad51 positive cells in 3 of the 4 PEO1-OR clones compared to PEO1 parental cells (Sup. Fig. 3C-D). Taken together, PARPi resistant cells have increased DNA damage repair capacity independent of *BRCA2* reversion mutations, which correlates with increased Wnt/β-catenin signaling.

Obtaining primary tumors of recurrent PARPi resistant disease is difficult given the natural progression of treatment (2). Therefore, we utilized a patient derived xenograft model (PDX) of a HGSOC primary tumor to recapitulate a recurrent tumor. Ascites containing malignant cells from a HGSOC patient were IP injected into nude SCID gamma mice and tumor bearing mice were treated with vehicle control or olaparib (50mg/kg daily). Following the cessation of olaparib, tumors were allowed to grow. After two months, ascites and tumor tissue were isolated from olaparib and control mice. RNA was extracted from ascites-derived tumor cells and *FOSL1* expression was assessed as an indicator of Wnt activation. In two of the four olaparib treated mice there was a significant increase in *FOSL1* expression compared to treated tumors suggesting a correlation between increased Wnt activation and olaparib insensitivity (Fig. 5A).

**Figure 5.**
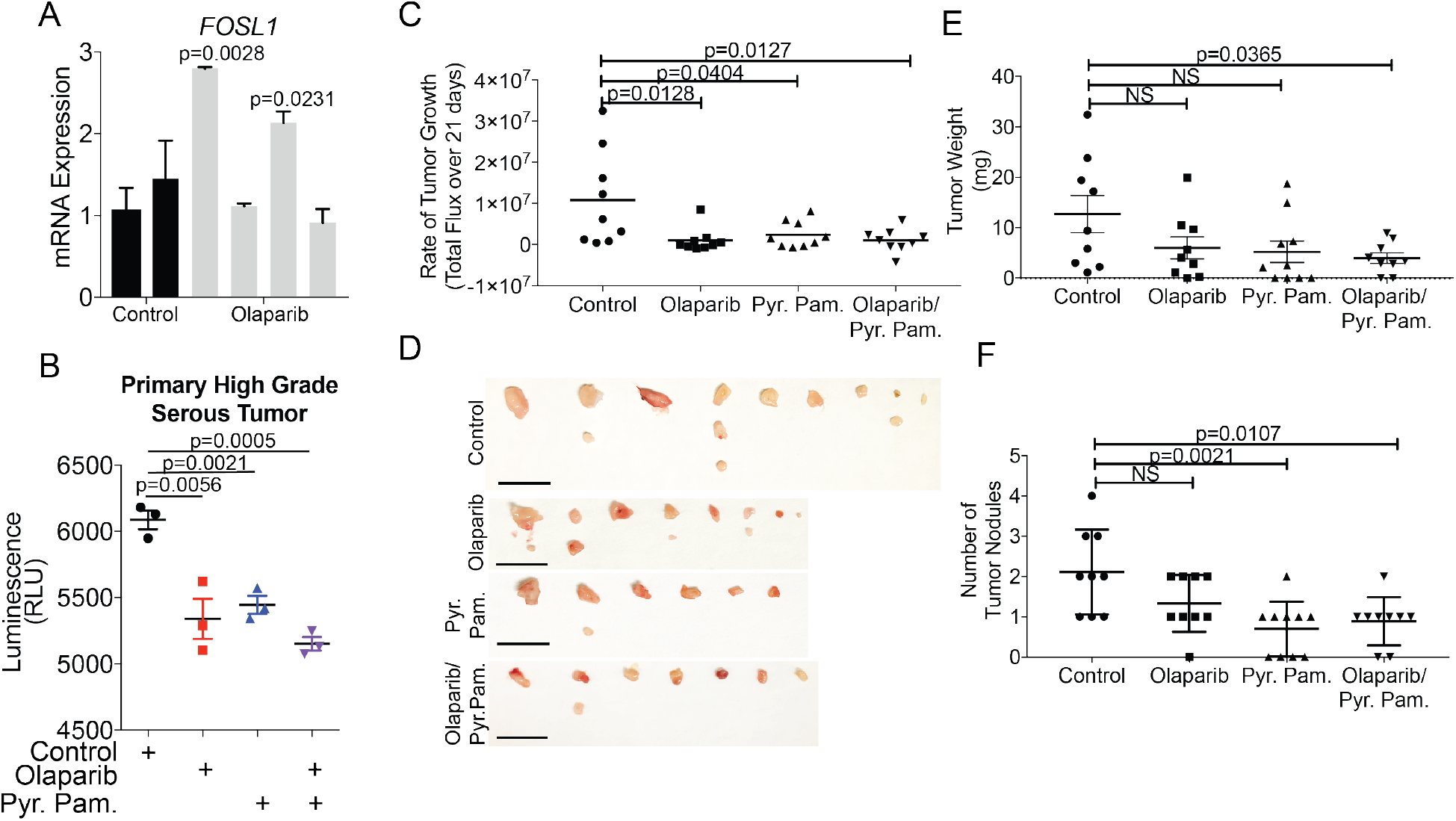
Combination of olaparib and Pyr. Pam. significantly inhibited HGSOC tumor progression. **A**) Primary HGSOC cells isolated from a patient were i.p. injected into immunocompromised mice. Tumor bearing mice were treated with vehicle control (control) or olaparib (50mg/kg, daily) for 21 days. Mice were monitored for 21days. Tumors and ascites were collected, RNA was extracted, and qRT-PCR was performed against *FOSL1. 18s* used as internal control. Each bar represents a different mouse. **B**) A primary high grade serous ovarian cancer was utilized for ex vivo culture. Tumor sections were tagged with a secreted luciferase and subsequently treated with olaparib and Pyr. Pam. for 48 hours. Cell culture media was refreshed and luciferase activity was measured after 4 hours. **C**) GFP/luciferase PEO1-WNT3A cells were injected into the intraperitoneal cavity of immunocompromised mice. Tumors were allowed to establish for 4 weeks. Mice were imaged and were randomized based in luminescence intensity on Day 0. Mice were treated daily for 21 days with vehicle control (n=9), olaparib (n= 9, 50mg/kg), Pyr. Pam. (n= 10, 0.5mg/kg) and olaparib/Pyr. Pam. (n=9, 0.5mg/kg). Mice were imaged twice a week. Mice were imaged and sacrificed on Day 22. Rate of luminescence change over the course of treatment graphed. **D**) Images of tumors derived from all groups. Scale bar = 1 cm. **E**) The weight of total tumor burden indicated as tumor weight. **F**) Number of tumor nodules from each group. Statistical used to calculate p-values, Analysis of variance (ANOVA). Error bars = SEM.

Utilizing an *ex vivo* model of a primary chemonaïve tumor we next examined the response of olaparib and/or Pyr. Pam. Primary tumors were sectioned with a Krumdieck tissue slicer, which produces uniform tissue slices that can be utilized for short-term culture. A HGSOC primary tumor was sectioned and tagged with a secreted luciferase (guassia luciferase; gLuc). gLuc activity has been utilized to measure proliferation and gLuc activity is directly correlated to cell proliferation (25). GLuc-tagged tumor sections were treated with olaparib and/or Pyr. Pam.. Following 72 hours, media was refreshed, gLuc was allowed to accumulate for 4 hours and the amount of gLuc activity was measured (Fig. 5B). Olaparib and/or Pyr. Pam. treated tumor sections had significantly reduced gLuc activity compared to the vehicle control. Taken together these in vivo studies suggest that hyperactivation of Wnt leads to olaparib resistance and that Wnt inhibition alone or in combination are viable therapeutic approaches.

According to the American Cancer Society, 80% of all HGSOC are diagnosed at late stage indicating that tumor has disseminated to the peritoneal cavity (1). Therefore, we wanted to examine the anti-tumor properties of combining olaparib with Pyr. Pam. in an intraperitoneal model of ovarian cancer. The *in vitro* results suggests an increase in Wnt signaling contributes to PARPi sensitivity and DNA damage repair therefore we wanted to recapitulate the WNT3A-dependent PARPi resistance *in vivo*. Therefore, we injected luciferase/GFP-tagged PEO1-WNT3A expressing cells into the peritoneal cavity of immunocompromised mice. Tumors were allowed to establish for four weeks and were randomized based on luciferase activity and mouse weight. Mice were then treated daily for 21 days with vehicle control, olaparib, Pyr. Pam., or in combination. Measured by total flux (photons/sec), we observed that all treatments resulted in a significant reduction in tumor growth rate compared to control mice (Fig. 5C and Sup. Fig. 4A). Total tumor burden at the end of 21 days of treatment was measured and only the olaparib/Pyr. Pam. treated tumors resulted in a significant decrease in tumor weight compared to control (12.7 vs. 3.978mg, p=0.0365, Fig. 5D-E). Dissemination of tumors was measured by quantifying GFP-positive tumor nodules during necropsy. Only mice treated with Pyr. Pam. and olaparib/Pyr. Pam. demonstrated a significant reduction in total tumor nodules (Fig. 5F). With respect to toxicity, although not significant we did observe that mice in all of the treatment groups had reduced body weight (Sup. Fig. 4B). The *in vivo* response observed in this xenograft model of HGSOC suggests that targeting Wnt signaling could prove to be an effective next-line therapeutic option following resistance to PARPi.

## DISCUSSION

Utilization of PARP inhibitors in the clinic is continuing to expand, which highlights an urgent need to better understand mechanisms of resistance. In this report, we established that HGSOC continually exposed to olaparib display hyperactivation of Wnt signaling and increased TCF transcriptional activity. The olaparib resistant HGSOC cells demonstrated increased DNA repair capacity, independent of *BRCA2* reversion mutations and that pharmacologic inhibition of Wnt signaling attenuated the DNA damage repair response. Moreover, PARP and Wnt inhibition significantly inhibited tumor progression in a HGSOC xenograft model and ex vivo cultures.

Transcriptome analysis of PEO1-OR cells revealed several significantly enriched signaling pathways and transcription factors, which highlights there are potentially multiple contributing pathways in promoting and maintaining PARPi resistance. We noted that PEO1 and OVCAR10 had an increase in TCF transcriptional activation, but OVSAHO cells did not have increased TCF activation. In the PEO1-OR TCF3 and LEF1 transcription factors were predicted to regulate a significant proportion of the olaparib resistant-related genes. Also, in a PDX model of recurrent olaparib insensitive tumors, a TCF target gene (*FOSL1*) was significantly upregulated. TCF/LEF transcriptional activation are highly dependent on Wnt/β-catenin activation. Therefore, we chose to further examine the relationship between Wnt signaling and PARPi response; however future work will evaluate the mechanism of olaparib resistance in OVSAHO cells. Wnt activation has been attributed to drive chemotherapy resistance in a variety of cancer types including prostate, colorectal, and ovarian (26–28). Wnt signaling is a driver of several major biological processes including proliferation, stemness, epithelial-mesenchymal transition, and DNA damage.

Beyond TCF/LEF transcription factors, we noted in PEO1-OR cells that the heterodimeric AP-1 transcription factor was identified as a top transcriptional regulator of differentially expressed genes. There is significant crosstalk between Wnt signaling and AP-1 regulation. Wnt signaling mediated TCF activity directly promotes expression of AP-1 subunits (*FOSL1, JUN*). Notably, p53, BRCA, NFκB, and AP-1 play critical roles in promoting the expression of DNA repair genes such as *ERCC4, ERCC6*, and *MGMT* [Reviewed in (29)]. HGSOC are often characterized by inactivation of TP53 and BRCA, therefore these cancers are potentially more dependent on AP-1 for transcriptional regulation of DNA damage repair effectors. Taken together, therapeutically targeting AP-1 could possibly provide a synthetic lethality in PARPi resistant HGSOC. Future studies will investigate the relationship between Wnt signaling, AP-1 activation, and PARPi resistance.

In this report, several Wnt inhibitors were examined, all of which have distinct mechanisms of action. Inhibitors of porcupine-mediated Wnt ligand secretion and β-catenin/CBP interaction failed to induce cytotoxicity effects in the HGSOC cell lines tested. In contrast, HGSOC cells were acutely sensitive to Pyr. Pam. induced β-catenin degradation, suggesting that future studies should focus on β-catenin stability. With respect to translatability, clinical studies have found that oral administration of Pyr. Pam. has relatively poor pharmacodynamics properties and limited bioavailability (30). Therefore, Pyr. Pam. in combination with olaparib was utilized as a proof-of-concept approach to examine tumor growth *in vivo*. Moreover, Pyr. Pam. significantly reduced TCF transcriptional activity indicating that an inhibitor of the β-catenin and TCF interaction could be an effective strategy to treat PARPi resistant HGSOC.

BRCA-reversion mutations are the most well-described adaptation that leads to PARP inhibitor resistance (31,32). Transcript and protein analysis did not reveal secondary BRCA2 reversion events in PEO1-OR cells. Independent of BRCA-reversion mutations, replication fork stability has been shown to promote PARPi resistance; however we did not find that the hyperactivation of Wnt signaling induced a robust change in RF stability. We did observe that HR and distal NHEJ DNA damage repair were increased in PEO1-OR cells compared to sensitive and that mhNHEJ seemed to be less important. Distal NHEJ and mhNHEJ are predominantly performed by a similar set of effector proteins, but are distinguished by the DNA polymerases, DNA protein kinase (DNA-PK) and nucleases [Reviewed in (33)], indicating that targeting the distal NHEJ pathway in PARPi resistant tumors is possible without inhibiting mhNHEJ.

In conclusion, hyperactivation of Wnt signaling contributes to and partially drives PARPi resistance independent of BRCA-reversion mutations. Inhibition of Wnt signaling reduces DNA repair capacity and significantly inhibits tumor progression *in vivo*. These findings offer a strong rationale to further examine the role of Wnt signaling in PARP inhibitor response.

## MATERIALS AND METHODS

### Cell lines and culture conditions

Epithelial ovarian cancer (EOC) cell lines were cultured in RPMI 1640 supplemented with 10% fetal bovine serum (FBS) and 1% penicillin/streptomycin. EOC cell lines (PEO1, OVCAR10, TOV-21G, and OVSAHO) were obtained from the Gynecologic Tumor and Fluid Bank (GTFB) at the University of Colorado. Viral packaging cells (293FT) were cultured in DMEM supplemented with 10% FBS at 37°C supplied with 5% CO_2_. Cells lines are authenticated at The University of Arizona Genomics Core using short tandem repeat DNA profiling. Regular Mycoplasma testing was performed using MycoLookOut Mycoplasma PCR detection (Sigma).

### Gynecologic Tissue and Fluid Bank (GTFB)

The University of Colorado has an Institutional Review Board approved protocol (COMIRB #07-935) in place to collect tissue from gynecologic patients with both malignant and benign disease processes. All participants are counseled regarding the potential uses of their tissue and sign a consent form approved by the Colorado Multiple Institutional Review Board. The tissues are processed, aliquoted, and stored at −80C.

### Retrovirus and lentivirus transduction

Retrovirus production and transduction were performed as described previously (34). Lentivirus was packaged using the Virapower Kit from Life Technologies (Carlsbad, CA) following the manufacturer’s instructions as described previously (9). shRNA and ORF were obtained from the Functional Genomics Facility at the University of Colorado. Cells transduced with virus encoding puromycin resistance gene were selected in 1 μg/ml puromycin.

### Reverse-transcriptase quantitative PCR (RT-qPCR)

RNA was isolated from cells with the RNeasy Mini Kit followed by on-column DNase digest (Qiagen). mRNA expression was determined using SYBR green Luna Universal One-step RT-PCR kit (New England Biolabs) with a BioRad CFX96 thermocycler. β-2-microglobulin (B2M), Glyceraldehyde 3-phosphate dehydrogenase (*GAPDH*), and 18s were used as internal controls. All primer sequences are in supplementary table 3.

### TCF transcriptional reporter assay

Using FuGENE6 transfection reagent (Promega) cells were transfected with either TOP-FLASH or FOP-FLASH. M50 Super 8x TOPFlash and FOPFlash were gifts from Randall Moon (Addgene *#* 12456 and 12457). Cells were incubated for 72 hours and subsequently lysed and analyzed using Luciferase Assay Kit (Promega), and luminescence was measured with a Promega GloMax.

### Colony formation assay

Cell lines were seeded in 24-well plates and treated with increasing doses of olaparib. Cell medium and olaparib were changed every two days with appropriate drug doses for 12 days. Colonies were washed twice with PBS, then incubated for 10 minutes in fixative (10% methanol and 10% acetic acid in PBS). Fixed colonies were stained with 0.4% crystal violet and rinsed in distilled water. Crystal violet was dissolved in fixative and absorbance was measured at 570nm.

### Reagents and antibodies

Olaparib, Rucaparib, WNT-C59, PRI724 were obtained from Selleckchem. Pyrvinium pamoate was obtained from Sigma Aldrich. The following antibodies were obtained from the indicated suppliers: BRCA2 (Bethyl, Cat#A303-434A, 1:2000), Vinculin (Cell Signaling Technology, Cat#13901, 1:1000), Wnt-3a (R&D systems, Cat# MAB9025-100, 1:1000), γH2Ax (Ser139) (EMD Millipore, Cat# 05-636, 1:1000), mouse anti-β-Actin (Abcam, Cat# ab6276, 1:10,000), Rabbit anti-β-Actin (Abcam, Cat# ab8227, 1:10,000), Rad51 (Abcam, Cat# ab176458). Anti-rat HRP (Jackson ImmunoResearch, Cat# 112-035-062, 1:5000)

### Immunoblotting

Protein was extracted with RIPA buffer (150mM NaCl, 1% TritonX-100, 0.5% sodium deoxycholate, 0.1% SDS, 50mM Tris pH 8.0) supplemented with Complete EDTA-free protease inhibitors (Roche), NAF and Na_3_VO_4_. Protein was separated on a SDS-PAGE and transferred to PVDF membrane using wet transfer or TransBlot Turbo (BioRad). Primary antibody incubation was performed overnight at 4°C. Secondary goat anti-rabbit (IRDye 680RD or IRDye 800CW, LI-COR, Cat # 92568071 or Cat # 926-32211, 1:20,000) and goat anti-mouse (IRDye 680RD or IRDye 800CW, LI-COR, Cat # 926-68070 or Cat# 925-32210, 1:20,000) antibodies were applied for one hour at room temperature. Bands were visualized using the Licor Odyssey Imaging System. For Wnt3a immunoblotting, antibodies were diluted in 5% milk/TBST (50 mM Tris pH7.5, 150 mM NaCl, 0.1% Tween20) blocking buffer and HRP chemiluminescent signal was detected with SuperSignal West Femto (Thermo scientific) and visualized using the G:Box (SYNGENE).

### RNA-Sequencing

RNA was isolated from PEO1 olaparib sensitive (n=2) and four PEO1 olaparib resistant clones using RNeasy columns with on-column DNase digest (Qiagen). RNA quality was confirmed using an Agilent Tapestation and all RNA used for library preparation had a RIN>9. Libraries were created using Illumina TruSEQ stranded mRNA library prep (#RS-122-2102). Strand-specific pair-ended Libraries were pooled and run on HiSeq4000 (Illumina). Library creation and sequencing were performed at the Genomics Core at the University of Colorado Anschutz Medical Campus. HISAT2 (35) was used for alignment against GRCh37 version of the human genome. Samples were normalized using TPM (Transcripts per Million) measurement and gene expression using the GRCh37 gene annotation was calculated using home-made scripts. Analysis was performed by the Division of Translational Bioinformatics and Cancer Systems Biology at the University of Colorado School of Medicine. GSE*####*

### Two-plasmid functional DNA repair assay

Two-plasmid functional assays were performed to assess distal non-homologous end joining (distal NHEJ), microhomology end-joining (mh-NHEJ), and homology directed repair (HDR). Cells were transfected with pimEJ5GFP (distal NHEJ) or EJ2GFP (mh-NHEJ) or pDRGFP (HDR). These plasmids all contain a mutated *GFP* reading frame. Transfected cells were selected with 1μg/mL puromycin and stably grown in 0.5μg/mL puromycin. Subsequently puromycin was removed and cells were transfected with I-SceI restriction enzyme. I-SceI introduces DNA double-strand breaks in the pimEJ5GFP (distal NHEJ) or EJ2GFP (mh-NHEJ) or pDRGFP (HDR) plasmids. Repair of the DNA damage through the indicated pathway results in restoration of a wildtype *GFP* reading frame and expression of GFP. After 72 hours post-I-SceI transfection, cells were collected and examined via a flow cytometer to quantify GFP positive cells. pimEJ5GFP and EJ2GFP were gifts from Jeremy Stark (Addgene plasmids # 44026 and # 44025). pDRGFP and pCBASceI were gifts from Maria Jasin (Addgene plasmid # 26475 and #26477).

### γ H2Ax resolution assay

Cells were plated in a 6-well plate (400,000/well) and irradiated with 5Gy. Irradiated cells were collected over a time course and utilized for protein extraction. Protein was used for immunoblot against γH2Ax and β-actin. Immunoblot signal was visualized on a Licor Odyssey and fluorescence signal was measured with the ImageStudio v4.0 software. γH2Ax signal was normalized to β-actin, graphed in Prism Graphpad, and followed by a linear regression analysis.

### Cell Viability Assay

HGSOC cells were seeded in a 96-well plate and treated for 72 hours. Cells were washed with PBS and incubated with MTT ((3-(4,5-Dimethylthiazol-2-yl)-2,5-Diphenyltetrazolium Bromide, Sigma Aldrich, St. Louis, MO) for 4 hours. Formazan was dissolved in DMSO and absorbance (590nm and 620nm) was measured on a Molecular Devices SpectraMax M2 microplate reader.

### Annexin V/Propidium Iodide assay

Phosphatidylserine externalization was detected using an Annexin V/propidium iodide (PI) staining kit (Life Technologies) following the manufacturer’s instructions. Annexin V/PI positive cells were detected using a Gallios Flow Cytometer (Flow Cytometry Core, University of Colorado) and analyzed with FlowJo software module.

### Animal Models

The protocols were approved by the <u>I</u>nstitutional <u>A</u>nimal <u>C</u>are and <u>U</u>se <u>C</u>ommittee (IACUC). For the recurrent olaparib-insensitive HGSOC model, patient-derived ascites (2.9 x 10^6^ cells) were intraperitoneal injected (COMIRB #07-935). After 7 days tumor bearing mice were randomized and treated with vehicle control (10% cyclodextrin) or olaparib (50/mg/kg, daily) for 21 days. Treated mice were monitored for 2 months. Tumor and ascites were collected and used for analysis. For olaparib and Pyr. Pam. combination experiment, the sample size of 9 mice per group was determined based on the data shown from *in vitro* experiments. Intraperitoneal xenograft was performed as described previously (36). Lentiviral particles specific for GFP/Luciferase were a gift from Diana Cittelly (37). GFP/Luciferase was transduced into PEO1-WNT3A cells and GFP positive cells were sorted twice. Briefly, 3.6 × 10^6^ luciferase-expressing PEO1-WNT3A cells were injected into the peritoneal cavity of 6-8-week-old female immunocompromised mice (n= 9-10 per group). PEO1-WNT3A cells were utilized for the *in vivo* evaluation because PEO1-OR cells failed to form IP tumors. 4 weeks after injection, tumors were visualized by injecting luciferin (i.p.: 4 mg/mice) resuspended in sterile PBS and imaged with an *In Vivo* Imaging System (IVIS). The mice were then randomized into 4 groups based on luciferase activity and mouse weight. Mice were treated with vehicle control (10% cyclodextrin), olaparib (daily, 50mg/kg), pyrvinium pamoate (daily, 0.5mg/kg), and with olaparib/pyrvinium pamoate. Mice were IP injected daily for 21 days and were imaged twice a week. Images were analyzed using Live Imaging 4.0 software. At end of the experiments, tumors were surgically dissected and tumor burden was calculated based on tumor weight. Peritoneal tumor nodules were quantified.

### *Ex vivo* Cultures

Fresh primary HGSOC tumor was obtained from the Gynecologic and Tumor Fluid Bank at the University of Colorado (COMIRB #07-935). Primary tumor was sectioned with Krumdieck Tissue Slicer to obtain 300 micron sections. Tumor sections were cultured with pBABE-puro-gLuc retrovirus for 24 hours. gLuc-tagged tissue sections were treated with olaparib (1μM), Pyr Pam (1μM) or in combination. Following a 72 hour incubation, gLuc activity was measured with Gaussia Luciferase kit (NEB) and analyzed on a Promega GloMax reader.

### Statistical Analysis

Statistical analyses and p value calculations were performed using GraphPad Prism 7 (GraphPad) for Mac OS. Quantitative data are expressed as mean ± SEM unless otherwise stated. Analysis of variance (ANOVA) with Fisher’s Least Significant Difference (LSD) was used to identify significant differences in multiple comparisons. Combination index was calculated using CompuSyn. For all statistical analyses, the level of significance was set at 0.05.

## ACKNOWLEDGEMENT

B.G. Bitler is supported by a NIH/NCI grant (R00CA194318). This work was supported in part by the University of Colorado Cancer Center’s Genomics and Microarray Core Shared Resource funded by NCI grant P30CA046934. Supported by NIH/NCATS Colorado CTSA Grant Number UL1 TR002535. Contents are the authors’ sole responsibility and do not necessarily represent official NIH views.

